# A novel xenograft model of human HCC in immunocompetent mouse

**DOI:** 10.1101/762351

**Authors:** Yanzhen Bi, Jun Shi, Shanshan Li, Quanyi Wang, Quanquan Wang, Xinyuan Wen, Feng Yang, Zhongping Duan, Yonghong Yang, Xiaobei Zhang, Yongchao Zhang, Zhenfeng Shu, Yu Chen, Feng Hong

**Author notes:** Corresponding authors: Feng Hong Institute of Liver Diseases, Affiliated Hospital of Jining Medical University, Jining, 272000, PR China., Yu Chen Difficult & complicated liver diseases and artificial liver center, Beijing Youan Hospital, Capital Medical University, Beijing, 100069, PR China. These authors contributed equally to this study (Yanzhen Bi, Jun Shi, Shanshan Li).

## Abstract

Hepatocellular carcinoma (HCC) is one of the most common malignant tumors that threaten human health; thus, the establishment of an animal model with clinical features similar to human liver cancer is of important practical significance. Taking advantage of the novel microcarrier-6, human HCC cells was injected into immunocompetent mice to establish a novel human HCC patient-derived xenograft (PDX) model. Primary HCC cells were isolated from fresh liver cancer tissues, which were subsequently co-cultured with microcarrier-6 to construct a three-dimensional tumor cell culture model *in vitro*. The HCC-microcarrier complex was implanted into mice by subcutaneous inoculation, and the tumor formation time, tumor formation rate, and pathological manifestation were recorded. Changes of immune parameters in mice were detected by flow cytometry. The success rate was 60% (6/10) in the establishment of liver cancer PDX mouse model, and the total tumor formation rate of the tumor-forming model is 80-100%. H&E staining and immunohistochemical experiments indicate that the model well retained the characteristics of the primary tumor. Interestingly, M2 macrophages in tumor-bearing mice increased significantly, and the levels of CD4^+^ T cells were significantly reduced. Through the application of the microcarrier-6 in immunocompetent mice, we successfully established a novel human HCC PDX model, which can be used to better study and further elucidate the occurrence and pathogenic mechanism of HCC.

## INTRODUCTION

Hepatocellular carcinoma (HCC), better known as liver cancer, is one of the most common malignant tumors and ranks in the top five among all cancers in terms of mortality rate (Bray et al., 2018; Ong et al., 2013). Early diagnosis of HCC remains difficult (Wang et al., 2016a), and the five-year survival rate is only 10-20% (Li and Wang, 2016; Liu et al., 2015); therefore, the establishment of an animal model of HCC that is capable of simulating these clinical features is of great significance for studying HCC. Human HCC patient-derived xenograft (PDX) animal models are based on the orthotopic or ectopic transplantation of fresh tumor tissue or tumor cells from patients into mice, providing a resource for tumor development (Kopetz et al., 2012). The PDX model is currently considered the most compatible animal model with clinical disease, and using fresh clinical HCC tissues can accurately reflect the HCC pathogenesis in patients (Bissig-Choisat et al., 2016; Hidalgo et al., 2014; Nicolle et al., 2016). However, the PDX model is typically constructed in highly immunodeficient mice to minimize the immune response, which imposes certain limitations to the study of tumorigenic mechanisms and the screening of antitumor drugs, in particular those dependent on the immune system. Therefore, an HCC PDX model in immunocompetent mice would be of great significance for the improvement of HCC treatment accuracy.

Microcarrier-6 is a novel microcarrier composed of a positively chargeable organic composite polymer. It has an irregular “maze”-like structure with multi-layered holes and strips to provide sufficient space. The pore size, surface positive charge density, and carrier particle size can be adjusted by chemical synthesis. It is a pure organic compound with advantages such as high purity, low immunogenicity, high biocompatibility, and being metabolizable and non-contaminating; thus, it can be used to provide a stable microenvironment for cell growth. The sufficient space available inside the microcarrier-6 solves the problems of uneven distribution of required nutrients and the produced metabolic waste. Simultaneously, due to its irregular “maze”-like structure, the microcarrier-6 can act as a barrier to prevent the direct attack of tumor cells by immune cells to a certain extent. In addition, following modification with stromal cell-derived factor-1α (SDF-1α) and vascular endothelial growth factor (VEGF), the microcarrier-6 can be directly embedded under the skin of mice, which accelerates blood vessel formation, induces blood vessel ingrowth, and provides a blood supply for rapid tumor growth.

In the present study, primary HCC cells were isolated from fresh liver cancer tissues and co-cultured with the microcarrier-6. The HCC-microcarrier complex was subsequently implanted into mice, resulting in the establishment of a novel HCC PDX model in immunocompetent mice. Moreover, we detected the changes of immune parameters of tumor-bearing mice, and the results showed that the M2-like macrophages promoting the tumor formation were activated, while the CD4^+^ T cells exerting the immune-clearing were suppressed.

## RESULTS

### The construction of a 3D human primary HCC cell culture system using the microcarrier-6

Microscopically, the isolated primary HCC cells were spherical, and almost all of them were single cells with a strong refractive index (Fig. 2A). The microcarrier-6 was observed as irregular lumps or long fusiform strips with a loose texture similar to fibrous tissue (Fig. 2B). Following a 24-h incubation of the primary HCC cells in medium, many small cell clusters were observed by microscopy, but single cells still accounted for the majority (Fig. 2C). Following 24 hours of co-culture, the primary HCC cells were firmly attached to the microcarrier-6 and reached a saturation state, with microcarriers surrounded by irregular cell clusters (Fig. 2D).

**Figure 1.**
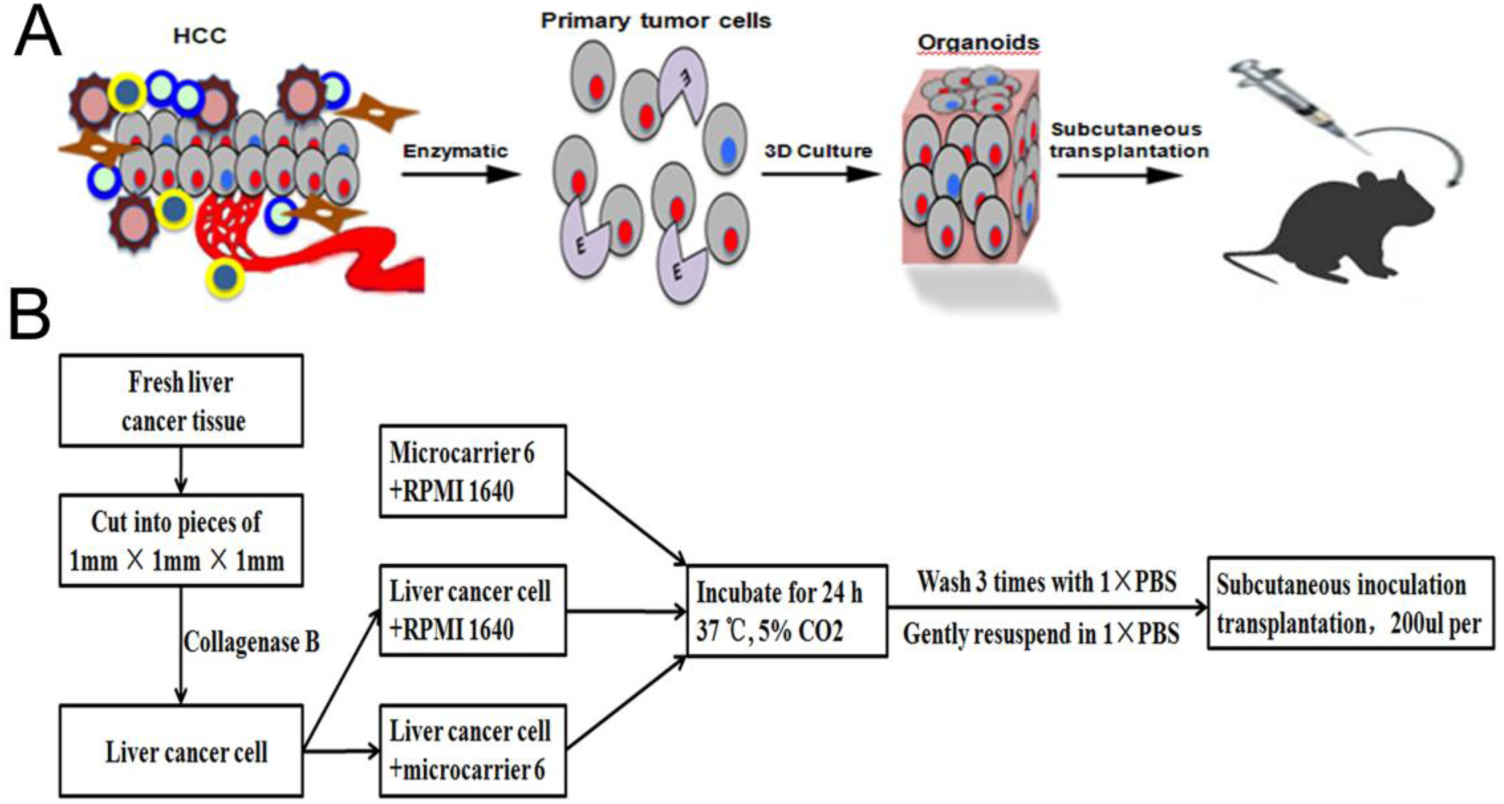
Schematic outline of modeling. A. Schematic outline of establishing HCC PDX model in immunocompetent mice. B. Different interventions in each group.

**Figure 2.**
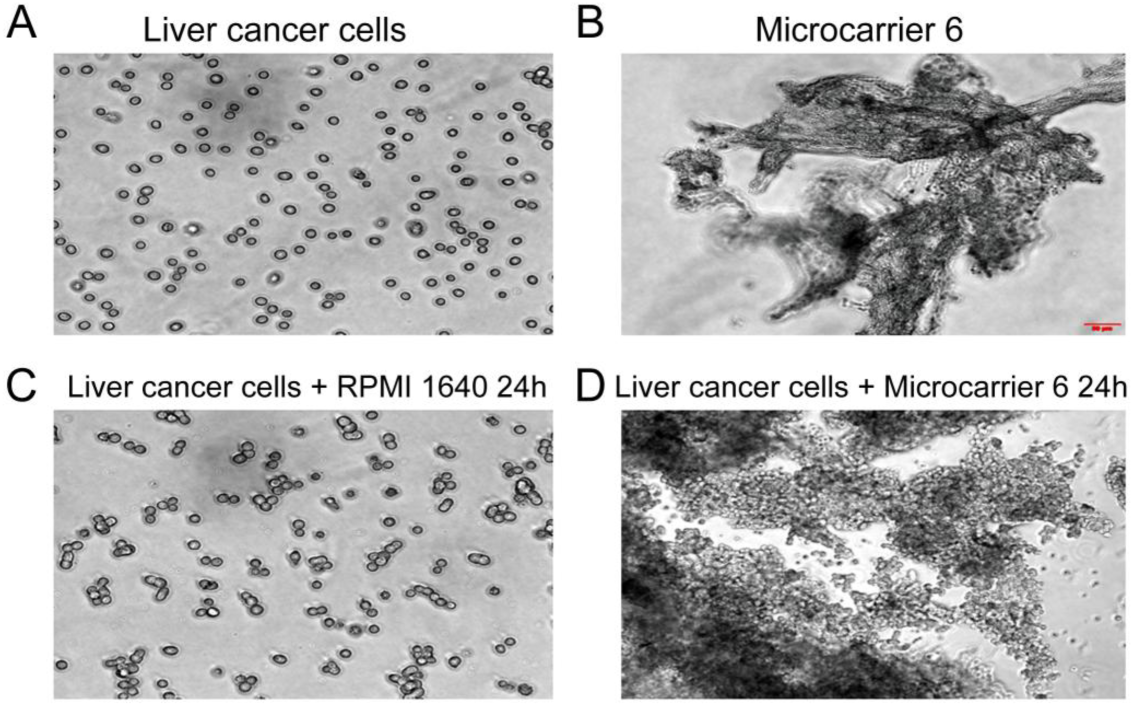
Three-dimensional co-culture system of liver cancer cells. A. Freshly isolated primary HCC cells were single cells with a strong refractive index. B. The microcarrier-6 was observed as irregular lumps or long fusiform by microscopy. C. Following a 24-h incubation of the primary HCC cells in medium, many small cell clusters were observed by microscopy. D. Following 24 hours of co-culture, the primary HCC cells were firmly attached to the microcarrier-6 scaffold.

### Establishment of an HCC PDX model in immunocompetent mice using the microcarrier-6

There was no significant change in appetite, coat, or weight of the mice in any group, and no mice died. Only the mice in the experimental group showed a slight decrease in activity after 1 week. No mice developed tumors in either the cell control group or the carrier control group during the course of the experiment (20 days) (Table 1, 2). While in the experimental group, 10 cases of liver cancer tissues were transplanted into immunocompetent mice, and finally 6 cases successfully constructed a novel xenograft model of human HCC (Table 1). The total tumor formation rate of the 6 tumor formation models was 90-100% (Table 2). In the experimental group, the transplanted tumors grew rapidly, and the subcutaneous mass was palpable 5 -7 days after inoculation (Fig. 3E). The peak growth period of the transplanted tumors lasted for 1-2 weeks, and a subcutaneous mass could be observed by the naked eye after approximately 2 weeks (Fig. 3E). At 2-3 weeks, the volume of the transplanted tumor changed little and reached the stationary phase. Simultaneously, the volume could reach 0.6 cm3 in approximately 20 days (Fig. 3C, 3D, 3E). The transplanted tumor tissue was easily separated from the surrounding tissue, and the shape was irregular, mostly round or elliptical. The surrounding blood supply was rich, displaying a grayish white and grayish red color (Fig. 3C, 3D).

**Table 1.** Summary of characteristics of the established PDX models for liver cancer.

| Case number | Tumor type | Gender | Age | Tumor diameter/cm | AFP/ug/L | Differentiation | Hepatitis | Cirrhosis or not | Strain | Implanted site | Tumor-formation or not |
| --- | --- | --- | --- | --- | --- | --- | --- | --- | --- | --- | --- |
| HCC-1 | Hepatocellular carcinoma | Female | 74 | 4.5×3.5×3 | 14.81 | Moderate | HBV | No | C57BL/6 | Subcutaneous | Yes |
| HCC-2 | Hepatocellular carcinoma | Male | 54 | 3×3×2.5 | 19.20 | Poor | HBV | Yes | C57BL/6 | Subcutaneous | Yes |
| HCC-3 | Hepatocellular carcinoma | Male | 46 | 13.5×12×8 | 58233.14 | Moderate | HBV | Yes | C57BL/6 | Subcutaneous | No |
| HCC-4 | Hepatocellular carcinoma | Male | 44 | 4.5×4×3.5 | 56.07 | Moderate or Poor | HBV | Yes | C57BL/6 | Subcutaneous | Yes |
| HCC-5 | Hepatocellular carcinoma | Male | 60 | 13×12×9 | 6424.04 | Moderate | HBV | Yes | C57BL/6 | Subcutaneous | No |
| HCC-6 | Hepatocellular carcinoma | Male | 64 | 4×3.5×3 | 7.7 | Moderate | HBV | Yes | C57BL/6 | Subcutaneous | No |
| HCC-7 | Hepatocellular carcinoma | Male | 61 | 5×4×4 | 372.55 | Moderate | HBV | Yes | C57BL/6 | Subcutaneous | Yes |
| HCC-8 | Hepatocellular carcinoma | Male | 47 | 5.5×5×4.9 | 1210.32 | Moderate or Poor | HCV | No | C57BL/6 | Subcutaneous | Yes |
| HCC-9 | Hepatocellular carcinoma | Male | 53 | 3×2×2 | 1524.85 | Moderate | HBV | No | C57BL/6 | Subcutaneous | Yes |
| HCC-10 | Hepatocellular carcinoma | Male | 58 | 7×4.5×4 | 1059.344 | Moderate | HBV | Yes | C57BL/6 | Subcutaneous | No |

**Table 2.** Tumor formation time and tumor formation rate.

| Groups | PDX-1<br>(HCC-1) | PDX-2<br>(HCC-2) | PDX-3<br>(HCC-4) | PDX-4<br>(HCC-7) | PDX-5<br>(HCC-8) | PDX-6<br>(HCC-9) | Tumor<br>formation<br>Time (day) | Total rate of<br>tumor<br>formation (%) |
| --- | --- | --- | --- | --- | --- | --- | --- | --- |
| Cell control<br>groups | 0/5 | 0/5 | 0/5 | 0/5 | 0/5 | 0/5 | — | — |
| Carrier control<br>groups | 0/5 | 0/5 | 0/5 | 0/5 | 0/5 | 0/5 | — | — |
| Experimental<br>groups | 4/5 | 5/5 | 4/5 | 4/5 | 5/5 | 5/5 | 5-7 | 80-100 |

**Figure 3.**
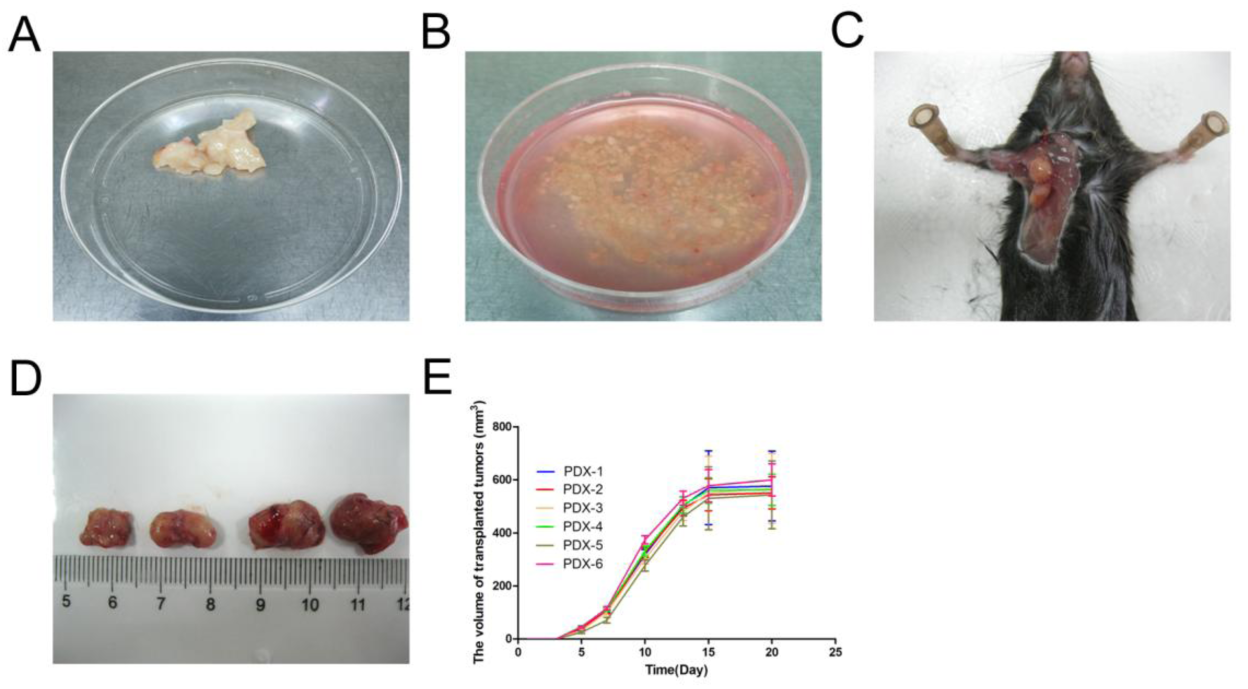
Characteristics of primary tumor and transplanted tumor tissue. A. Fresh primary liver cancer tissue. B. The primary HCC cells were separated from comminuted tumor tissues by an enzymatic method. C. Tumor-bearing mouse. D. Transplanted tumors. e Growth curves of transplanted tumors.

### H&E staining of the primary and transplanted tumors

Following application of H&E staining to the primary HCC tissue paraffin sections, numerous abnormally shaped and disordered tumor cells could be observed by light microscopy (Fig. 4A, 4B). Similarly, the transplanted tumor tissue also showed a large number of disordered (in a nest or flaky shape) and obviously atypical cells, with a round or elliptical shape. These cells had a large nucleus, coarse chromatin, and obvious nucleoli of different sizes, showing pathological mitotic characteristics. A large number of lymphocytes and a small amount of foreign matter around the microcarrier-6 that had not been cleared could be seen in the stroma. HCC cells had also infiltrated the peripheral fat and muscle tissues, accompanied by obvious necrosis mainly occurring in the center of the tumor. The newly developed capillaries in the transplanted tumor tissue were abundant, mostly located at the edge of the tumor (Fig. 4C, 4D).

**Figure 4.**
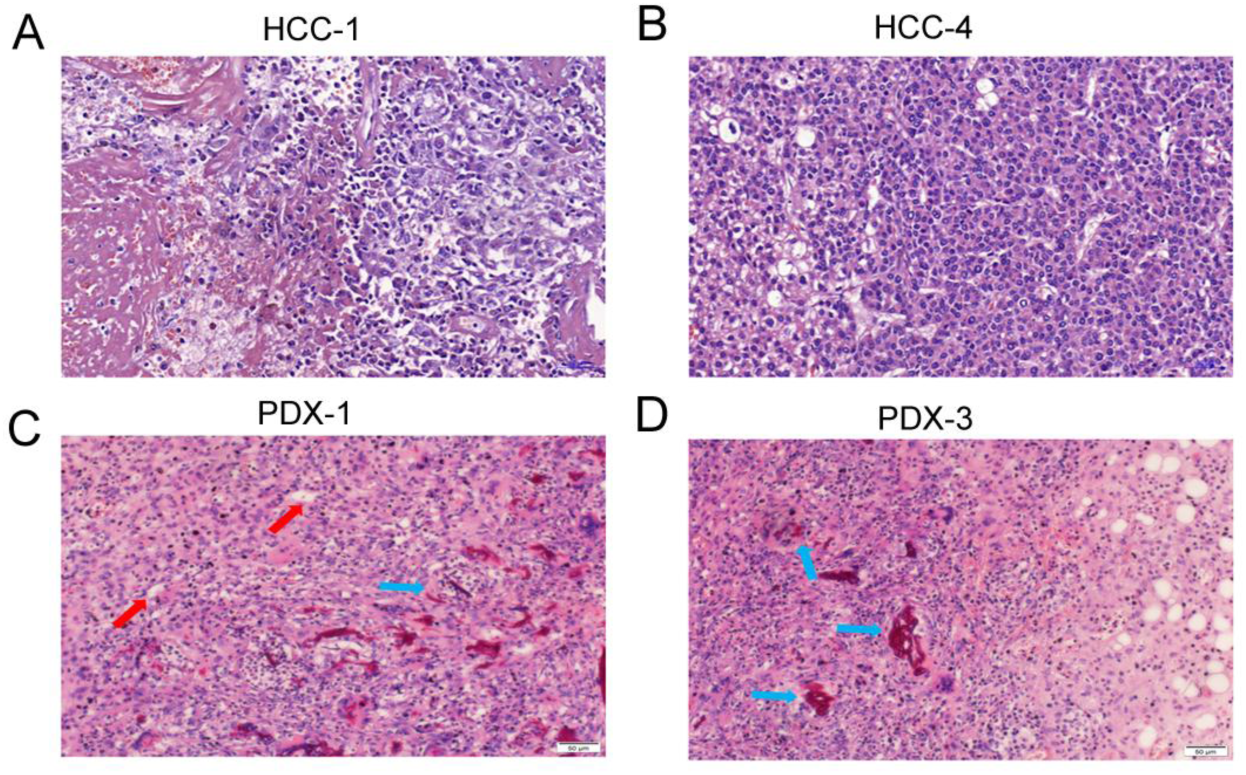
H&E staining of primary tumors (A, B) and transplanted tumors (C, D). A, B. Numerous abnormally shaped and disordered tumor cells could be observed by light microscopy in the primary HCC tissue. C, D. The transplanted tumor tissue also showed a large number of disordered and obviously atypical cells. A small number of microcarriers surrounded by inflammatory cells that form granulomas. Abundant vessels were shown in tumor tissues. Red arrows refer to blood vessels and blue arrows indicate granulomas.

### Immunohistochemical staining of the primary and transplanted tumors

Gpc-3, Hep-1, and CK8/18 are mainly expressed in the membrane and cytoplasm of human cancer cells, and therefore are specific markers for the human HCC cells. The immunohistochemical staining revealed that Gpc-3, Hep-1, and CK8/18 were all positively expressed in the primary (Table 3) and transplanted tumor tissues(Fig. 5), confirming that the atypical cells were human-derived HCC cells.

**Table 3.** Immunohistochemical staining of patient tumor tissues and patient-derived transplanted tumor tissues

| Case number | GPC-3 | Hep-1 | CK8/18 |
| --- | --- | --- | --- |
| HCC-1 | + | + | + |
| HCC-2 | + | + | + |
| HCC-3 | + | + | + |
| HCC-4 | + | + | + |
| HCC-5 | + | + | + |
| HCC-6 | + | + | + |
| HCC-7 | + | +/- | + |
| HCC-8 | + | + | + |
| HCC-9 | + | + | + |
| HCC-10 | + | + | + |
| PDX-1 | + | + | + |
| PDX-2 | + | + | + |
| PDX-3 | + | + | + |
| PDX-4 | + | + | + |
| PDX-5 | + | + | + |
| PDX-6 | + | + | + |

**Figure 5.**
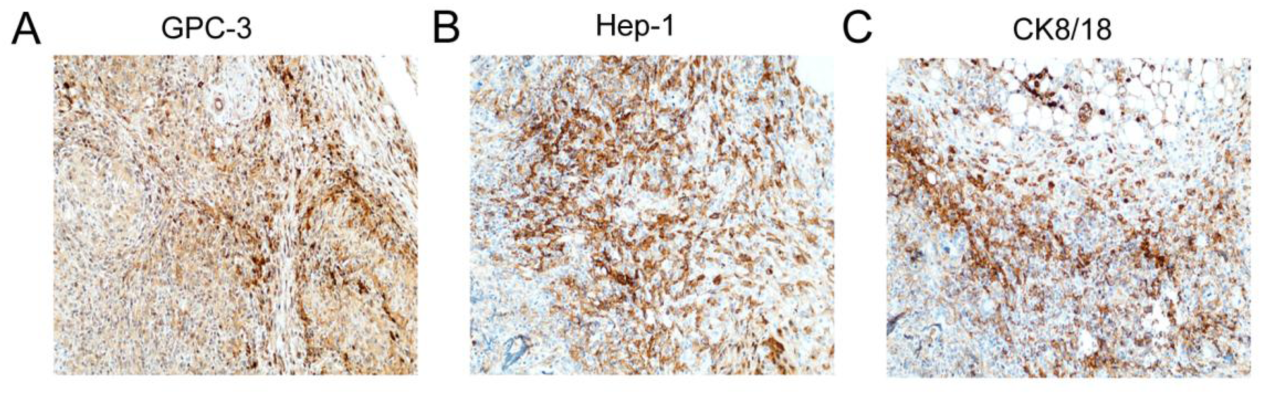
Immunohistochemical staining of transplanted tumors. The human liver cancer xenograft displaying diffuse and strong immunoreactivity in the cytoplasm of cancer cells. A. Gpc-3. B. Hep-1. C. CK8/18.

### Changes of macrophages and CD4^+^ T cells during tumor formation in mice

To evaluate the changes of immune cell subpopulations during tumor formation in mice, we performed flow cytometry analyses of macrophages and CD4^+^ T cells in spleen samples (Fig. 6). As shown in Fig. 6, tumor-bearing mice in experimental group (PDX) showed a significantly higher frequency of macrophages and decreased prevalence of CD4^+^ T cells in the spleen as compared with control mice. There was an increased prevalence of M2 macrophages in the spleen of tumor-bearing mice as compared with non-tumoid mice in experimental group (PDX-) (Fig. 6B), while the percentage of M1 macrophages and CD4^+^ T cells in the spleen was significantly decreased (Fig. 6A, 6C). The above-mentioned results indicate that the M2-like macrophages promoting the tumor formation were activated, while the CD4^+^ T cells exerting the immune-clearing were suppressed.

**Figure 6.**
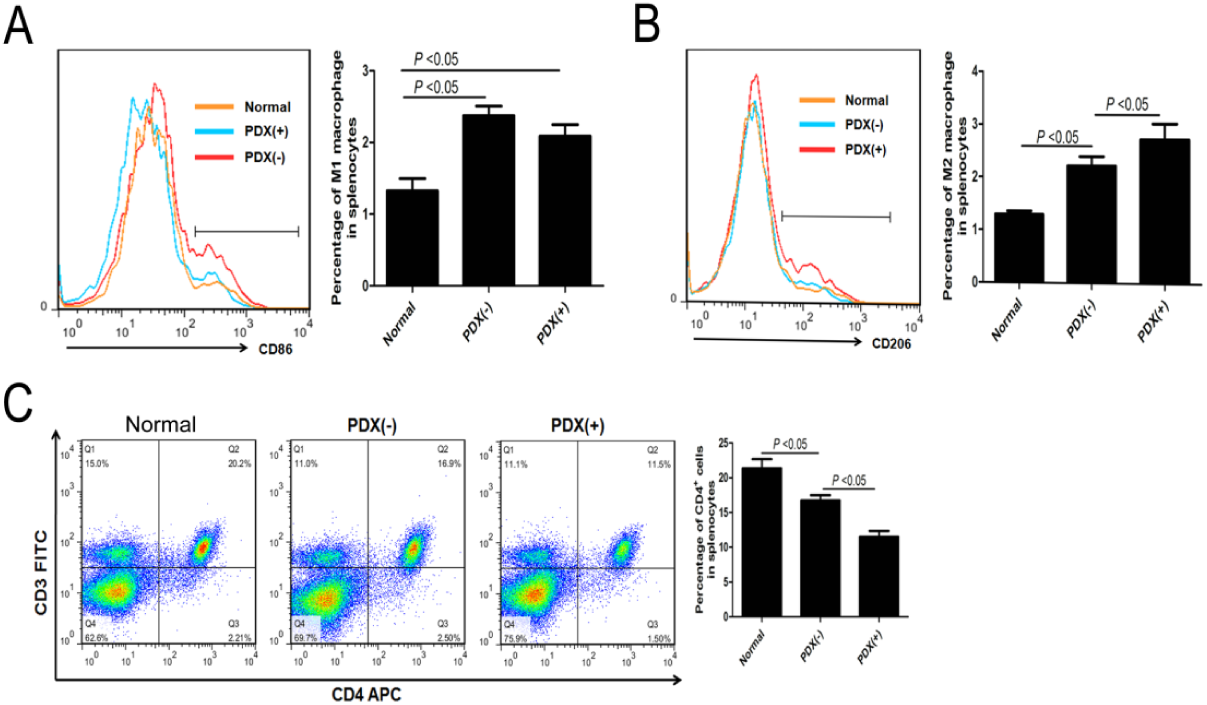
The changes of immune cell subpopulations during tumor formation in mice. The number of spleen macrophages (A, B) and spleen CD4^+^ T lymphocytes (C) in mice were analyzed by FACS at 10 days. PDX (+) represented tumor-bearing mice in the experimental group, while PDX (-) represented non-tumoid mice in the experimental group. Data are shown as the mean ± SEM, as determined by ANOVA; ns denotes *p* > 0.05.

### DISCUSSION

HCC is one of the most common malignant tumors that threaten human health. Currently, liver cancer is mainly treated using surgery, chemotherapy/radiotherapy combination treatment, and gene-targeted therapy; however, there is still no effective way to completely control the development of HCC. The discovery of novel treatments for HCC requires animal-based experiments, and the establishment of an animal model that mimics human liver cancer is therefore essential for HCC research. In recent years, with the advance of related technologies, comprehensive research has been conducted to develop novel HCC models, among which the HCC orthotopic transplantation models, the induced HCC models, the transgenic HCC models, and the human HCC xenograft models are the main ones.

The ideal HCC model should accurately reflect the biological characteristics of HCC, fully simulating the human tumor microenvironment. Moreover, it should be easy to operate and have a high reproducibility and a low price. To date, the HCC PDX model is a commonly used model with the most comparable clinical features to humans, and serves as an important platform for the study of preclinical evaluation, tumor treatments, and prognosis. It is expected that this model will result in novel breakthroughs in individualized treatments (Izumchenko et al., 2016; Siolas and Hannon, 2013; Tentler et al., 2012). The PDX models established thus far are usually in immunodeficient mice, which are not only expensive to raise, but also cannot be used for the evaluation of the efficacy of immune-related drugs (e.g., vaccines, immunomodulatory PD-1 antibody drugs, or anti-tumor drugs functioning by immune activation, such as the CD40 monoclonal antibodies) (Kim et al., 2009; Pavia-Jimenez et al., 2014; Zhang et al., 2013). In addition, the existing PDX model does not reflect the intrinsic link between tumor development and immune status/regulation.

In the present study, immunocompetent C57BL/6 mice were used to overcome the above problems. In addition, the pathological type of the ten clinical samples selected in this experiment was HCC, which is the most common type of primary liver cancer, and thus is clinically representative. Subcutaneous ectopic transplantation of the tumors under the armpit mainly considered the rich blood supply and tissue loosening in this site, which are conducive to tumor growth (Zheng et al., 2012).

The microcarrier-6 used in the present study is a novel type of microcarrier, which was co-cultured with human primary HCC cells to successfully construct a 3D growth model (Fig. 2D). Since the microcarrier-6 has a low immunogenicity, a loose texture, and a large number of central pores for the growth of tumor cells, it can act as a barrier to block the direct killing of tumor cells by immune cells. Following modification, the microcarrier-6 can more easily allow the growth of blood vessels, providing good conditions for the rapid growth of tumor cells. The 3D cell mass formed by the 24-h co-culture of tumor cells with the microcarrier-6 prevented single tumor cells from being rapidly cleared by the mice immune system. Moreover, this approach also effectively prevented cell death which plagued the interior of the transplanted primary tissue block due to lack of blood supply.

With the newly established 3D growth model of human primary HCC, we successfully developed a PDX mode of HCC in immunocompetent mice in the experimental group. The mice in the cell control and carrier control groups did not develop tumors. This PDX model is characterized by rapid tumor growth, which can be detected in 5-7 days and peaks at 1-2 weeks, reaching a volume of 0.6 cm3 in approximately 20 days. Cellular immunity is the main force against tumor growth; the cells involved mainly include T cells, natural killer (NK) cells, macrophages, and dendritic cells (DC) (Wang et al., 2016b; Zitvogel et al., 2016). Interestingly, we found that the M2-like macrophages promoting the tumor formation were activated, while the CD4^+^ T cells exerting the immune-clearing were suppressed during the early stage of transplanted tumor formation. This indicates that foreign tumor cells can regulate the immune system of mice and create a suitable microenvironment for the rapid growth of tumor cells.

Pathological H&E staining revealed a large number of cells with an atypical nucleus that had infiltrated into muscle, fat, and other tissues, in addition to a large area of necrosis in the center of the tumor, which is in accordance with the development of human HCC, considering the fast growth and insufficient blood supply. The capillaries, mainly located around the tumor, were also abundant. Immunohistochemistry of the liver cancer-specific markers Gpc-3, Hep-1 and CK8/18 were positive, confirming that the heteromorphic cells were human HCC cells.

Through the inoculation of human primary HCC cells into immunocompetent mice, the present study successfully established a human HCC PDX model using the microcarrier-6. This model can reflect the interaction between the immune system and tumors; therefore, it is anticipated to bring considerable broad benefits to the cancer research community.

## MATERIALS AND METHODS

### Reagents

Collagenase B was purchased from Sigma (Shanghai, China). RPMI 1640 medium, trypsin, fetal bovine serum, red blood cell lysate, and streptomycin were purchased from Gibco (Thermo Fisher Scientific, Waltham, MA). Rabbit anti-human CK8/18, GPC-3, and Hep-1 monoclonal antibodies were purchased from Abcam (Cambridge, UK), and isoflurane was purchased from Shenzhen, China. The microcarrier-6 was supplied by ELYON BIO-TECHNOLOGIES LLC (Gaithersburg, MD).

### Experimental animals

150 C57BL/6 mice (male, 6-8 weeks old, body weight 22-25 g) were purchased from Jinan Pengyue Experimental Animal Breeding Co., Ltd. [License No. SCXK 20140007]. The mice were raised in the animal center of the Affiliated Hospital of Jining Medical College (SPF class). All animal experiments were performed in accordance with the relevant regulations of the Animal Ethics Committee of the Affiliated Hospital of Jining Medical College.

### Patient information

Ten fresh HCC specimens were obtained from the Department of Hepatobiliary Surgery of the Affiliated Hospital of Jining Medical University. The acquisition of tumor specimens was approved by the patients and their family members, and the informed consent was signed, clarifying that the obtained specimens were for research purposes only. The present experiment was approved by the Ethics Committee of the Affiliated Hospital of Jining Medical University. The details of the ten liver cancer patients are shown in Table 1.

### Acquisition of human primary HCC cells

A collagenase B solution (0.05% in RPMI 1640) was filtered through a filter with a pore size of 0.22 μm, and placed in a thermostatic water bath at 37°C for 30 minutes. Fresh HCC specimens were washed three times in RPMI 1640 medium and cut into pieces (approximately 1 mm × 1 mm × 1 mm) with scissors (Fig. 3A, 3B). The specimen pieces were mixed with 0.05% collagenase B solution and incubated at 37°C for digestion. After 1 h, the mixture was diluted in RPMI 1640 and mixed using a pipette. The extracted supernatant was filtered through a sieve with a pore size of 70 μm, and centrifuged at 1000 rpm for 8 min. Following the lysis step of red blood cells, human primary HCC cells were obtained. Residual tissue blocks were mixed with 0.05% collagenase B for further digestion. Human primary HCC cells were sequentially extracted after 2, 3, and 4 h of collagenase B digestion, as described above (Fig. 1, 2A).

### Co-culture of human primary HCC cells with the microcarrier-6

The microcarrier-6 was immersed in 75% ethanol for 24 h, washed three times with 1 × phosphate -buffered saline (PBS), and added to RPMI 1640 medium containing 10% fetal bovine serum; the concentration of the microcarrier suspension was subsequently adjusted to 300 μg/mL. Following incubation for 24 h at 37°C, the microcarrier-6 was ready for use. Subsequently, the microcarrier-6 was incubated with SDF-1α and VEGF at a concentration of 100 ng/mL for 12 h. Human primary HCC cells extracted at all time points were pooled, resulting in a high percentage of viable cells, as counted by trypan blue staining (> 95%). The cell concentration was adjusted to 2 × 10^7^/mL by resuspending the cells in RPMI 1640 medium containing 10% fetal bovine serum. Half the liver cancer cell suspension was mixed with the modified microcarrier-6 suspension (1:1 v/v) in a 15-mL centrifuge tube, and the other half was added to an equal volume of RPMI 1640 medium containing 10% fetal bovine serum. Moreover, an equal volume of microcarrier-6 suspension was added to RPMI 1640 medium containing 10% fetal bovine serum. These three tubes were placed in an incubator at 37°C, 5% CO_2_ for 24 h (Fig. 1, 2).

### Establishment of an HCC PDX model in immunocompetent mice

The ten specimens were shared among the 150 C57BL/6 male mice, 15 in each case, which were subsequently sub-divided into three groups: cell control group, carrier control group, and experimental group, with five mice in each group. The mice in cell control groups were inoculated with HCC cell suspension only at 2 × 10^6^ cells/mouse (counted prior to the co-culture step); those in carrier control groups were inoculated with the microcarrier-6 suspension only at 30 μg/mouse; and those in experimental groups were inoculated with the HCC-microcarrier complex suspension at 2 × 10^6^ cells and 30 μg microcarrier-6 (counted prior to co-culture). Following a 24-h co-culture, all samples were washed three times in 1 × PBS, gently mixed with 1 × PBS to the same final volume as before washing, and placed on ice until further use. The mice were anesthetized with isoflurane and inoculated with each individual co-culture solution using a trocar in the right axilla (200 μL/mouse) (Fig. 1).

### Observation indicators and pathological examination

The longest (a) and shortest (b) diameters of the subcutaneously transplanted tumors were measured once a week, the tumor volume (Ogawa et al., 2007) was calculated using V = 1/2 × a × b^2^, and a tumor growth curve was constructed. When the tumors had a long diameter of 1 cm, the mice were sacrificed by cervical dislocation, and the tumor tissues were completely excised. Following fixation in 4% neutral formaldehyde and paraffin embedding, the tissues were stained with H&E. Immunohistochemical staining was performed using the EnVision two-step method according to the manufacturer’s instructions. The staining results were interpreted as: positive if the positive particles located in the tumor cytoplasm represented ≥ 5% of the total cells; 5%, negative.

### Flow cytometry

Spleen samples were collected and ground and filtered into a 15 mL centrifuge tube and centrifuged at 1200 rpm for 5 min. Spleen cells can be obtained after lysis of red blood cells. For phenotypic staining, cells were washed twice with PBS containing 1% FBS and 0.1% NaN3, and subsequently incubated with mouse primary antibodies against CD3, CD4, CD11b, F4-80, CD86 and CD206 for 30 min at 4°C according to the standard procedure. After washing twice with PBS, cells were analyzed using a FACSCalibur flow cytometer (BD Biosciences). An isotype control was used for each antibody.

### Statistical analysis

The SPSS 20.0 software (IBM SPSS, Chicago, IL) was used for statistical analyses. Measurement data are presented as the mean ± standard error of the mean (SEM). The differences among the mean values of each group were compared using the one-way analysis of variance (ANOVA) or Kruskal-Wallis test as appropriate. *p* < 0.05 is considered a statistically significant difference.

## Author contributions

Conceptualization: Y.C., F.H., Y.B.; Methodology: J.S., S.L, Q.Y.W., Q.Q.W.; Validation: X.W., F.Y.; Formal analysis: Z.D., Y.Y.; Investigation: X.Z., Y.Z.; Resources: X.W., F.Y., Z.S.; Data curation: F.H., Y.B., Q.Q.W., S.L.; Writing - original draft: Y.B.; Writing - review & editing: Y.C., F.H.; Visualization: Y.C., F.H., Y.B.; Supervision: F.H.; Project administration: F.H.; Funding acquisition: Y.C., F.H.

